# A novel *cis* regulatory element regulates human *XIST* in CTCF-dependent manner

**DOI:** 10.1101/871178

**Authors:** Rini Shah, Ankita Sharma, Ashwin Kelkar, Kundan Sengupta, Sanjeev Galande

**Affiliations:** 1Centre of Excellence in Epigenetics, Department of Biology, Indian Institute of Science Education and Research, Pune, India; 2Chromosome Biology Laboratory, Department of Biology, Indian Institute of Science Education and Research, Pune, India

**Keywords:** XIST, *cis* regulatory element, pluripotency factors, YY1, CTCF

## Abstract

The long non-coding RNA XIST is the master regulator for the process of X chromosome inactivation (XCI) in mammalian females. Here we report the existence of a hitherto uncharacterized *cis* regulatory element (cRE) within the first exon of human *XIST*, which determines the transcriptional status of *XIST* during the initiation and maintenance phases of XCI. In the initiation phase, pluripotency factors bind to this cRE and keep *XIST* repressed. In the maintenance phase of XCI, the cRE is enriched for CTCF which activates *XIST* transcription. By employing a CRISPR-dCas9-KRAB based interference strategy, we demonstrate that binding of CTCF to the newly identified cRE is critical for regulating *XIST* in a YY1-dependent manner. Collectively, our study uncovers the combinatorial effect of multiple transcriptional regulators influencing *XIST* expression during the initiation and maintenance phases of XCI.

## INTRODUCTION

X chromosome inactivation is a gene dosage compensatory phenomenon in the mammals. XIST, the inactive X (Xi)-specific transcript, is the long non-coding RNA that co-ordinates the process of X chromosome inactivation (XCI) in the eutherian mammalian females (1–3). The phenomenon of XCI corrects for the X-linked gene dosage disparity between the males (XY) and females (XX) of mammalian species (4). The first and the foremost event for the initiation of XCI is the mono-allelic and sustained upregulation of Xist which occurs in the epiblast cells during the implantation of the mouse embryos. At this stage, one of the two X chromosomes is randomly chosen for silencing (5, 6). Xist/XIST lncRNA physically coats the chosen Xi *in cis,* initiating epigenetic reprogramming characterized by the sequential removal of active chromatin marks, RNA Pol II exclusion and the establishment of repressive modifications (7–10), macroH2A recruitment (11) and CpG methylation of gene promoters (12, 13). Consequently, the entire X chromosome undergoes heterochromatinization, rendering it stably inactive in a mitotically heritable manner (14–22). Since Xist/XIST lncRNA is the central player for establishing the process of chromosome-wide transcriptional silencing, it is essential to regulate its levels and function to ensure proper initiation of XCI as well as maintenance.

Several studies have demonstrated that the temporal activation of *Xist* during a specific window of development is brought about by the concerted action of a network of activators and repressors either encoded from the X-inactivation centre (Xic) locus or regulating Xic (23). In addition to *Xist*, the Xic locus codes for a number of lncRNAs that participate in regulating *Xist*. One of the most critical *cis*-regulators of *Xist* is its antisense lncRNA–Tsix, which unlike Xist, is exclusively produced from the future active X (Xa) and negatively regulates *Xist* by modulating the chromatin architecture of its promoter (24–28). Xic is partitioned into two distinct topologically associated domains (TADs) – (i) *Xist* TAD (∼550 Kb) containing *Xist* and its positive regulators such as *Ftx* and *Jpx*, (ii) *Tsix* TAD (∼300 Kb) harbouring negative regulators of *Xist* such as *Tsix* and Xite. Both these TADs show opposite transcriptional behaviour on the chosen Xi, with the expression of genes on Xist TAD increasing and that on Tsix TAD decreasing during differentiation of mouse embryonic stem cells (mESCs) (29, 30). A recent report highlighted the role of promoter of another long non-coding RNA, Linx as a *cis*-regulatory silencer of *Xist*. *Linx* is located on Tsix TAD and serves as a silencer of Xist independent of its transcript or transcription or Tsix. However, when it is placed in *Xist* TAD, it serves as an enhancer of Xist (31). Besides the X-chromosomal *cis*-acting modulators, the autosomally encoded *trans*-acting factors such as the core pluripotency factors – Oct4, Sox2, Nanog, Klf4 and c-Myc link the two developmental events of differentiation and XCI (32), by regulating *Xist* (33, 34),*Tsix* (35, 36)*, Rnf12* (37) and *Linx* (31). Thus, it can be convincingly stated that Xist expression is robustly regulated by a multitude of *cis* and *trans* factors acting either synergistically or independently to ensure accurate execution of the developmentally important process of XCI.

For the past two decades, mouse has been the preferred model system to study the molecular pathways leading to the initiation and establishment of XCI. Our understanding of the mechanism(s) regulating *XIST* and the molecular dynamics of XCI in other eutherian mammals is rather limited. Deciphering the XCI pathways in multiple systems is important to address the question of conservation and evolution of the process of XCI. Although mouse and human *Xist*/*XIST* were discovered almost simultaneously (1–3), our understanding of human *XIST* regulation has remained poor as compared to its mouse counterpart. It has been demonstrated that human and mouse *XIST*/*Xist* are functionally conserved since ectopic insertion of human *XIST* in murine and human cells induces XCI (17, 18, 38, 39). However, there is only 49% conservation at the sequence level, with the maximum homology observed in the first exon which harbours repeat elements (A-F) (40–42). Most notably, *Tsix*, the key negative regulator of *Xist* in mouse is a pseudogene in humans and the potential regulatory sequences of *XIST* also do not show any conservation between humans and mouse (43–45). Hence, apart from a plethora of other factors regulating *Xist*, the critical mechanism governed by *Tsix* does not seem to be conserved. It is known that Xist/XIST is not only induced in both male as well as female blastocyst from the maternal as well as paternal X chromosomes, it also coats the X chromosomes, leading to partial silencing of X-linked genes in mouse but not in humans (46, 47). This discrepancy can be attributed to the human specific lncRNA, XACT, which specifically coats Xa and co-accumulates with XIST in human preimplantation as well human embryonic stem cells (hESCs), possibly tampering with XIST silencing function (48, 49). This suggests that although the process of XCI is conserved across eutherians and is dependent on XIST RNA, this process could be manifested in diverse ways in different species.

A few studies have attempted to address the regulation of human *XIST*. Hendrich *et al*. compared the putative promoter sequences from human, horse, rabbit and mouse and discovered that the first 100 base pairs (bp) upstream of the Transcription Start Site (TSS) exhibit maximum conservation and hence assigned it as the promoter of human *XIST* (50). Through a series of *in vitro* biochemical assays, the authors identified three transcription factors – SP1, YY1 and TBP as potential regulators of *XIST*. The question whether any of these factors can bind *XIST* promoter in cells and regulate its expression was addressed by two independent studies wherein YY1 was uncovered as the key transcription factor activating *XIST* transcription (51, 52). Here, we demonstrate for the first time that besides YY1, *XIST* transcription is also under the control of pluripotency factors – OCT4, SOX2 and NANOG as well as CTCF. More specifically, *XIST* is repressed by the pluripotency factors and activated by YY1 and CTCF. We report the existence of a novel *cis* regulatory element (cRE) at the *XIST* locus located in the first exon that seems to be bound by the aforementioned factors in a sex-specific and X inactivation-dependent manner. Further, this element could presumably act as a crucial determinant of transcriptional outcome from the *XIST* promoter during the initiation as well as maintenance phases of XCI.

## RESULTS

### Cell line models provide the context for initiation and maintenance phases of XCI

XCI can be broadly categorised into two stages, (i) Initiation - when differentiating embryonic stem cells undergo XCI for the first time and (ii) Maintenance - where any cell carrying an inactive X ensures continued expression of XIST RNA from the inactive X upon subsequent cell divisions. In the case of human XCI, YY1 is the only known regulatory factor that influences XIST expression from the inactive X. However, the question of alternate regulatory molecules affecting XIST expression is still unanswered. This is especially of importance in lieu of the fact that the antisense RNA *TSIX* is truncated/non-functional in humans (43, 44).

To study the regulation of *XIST* promoter-related factors other than YY1 in the contexts of initiation and maintenance phases of XCI, we employed a human male embryonic carcinoma cell line, NTERA-2 clone D1 (NT2/D1), and differentiated cells such as HEK293T respectively. NT2/D1 cells express pluripotency factors, low amounts of XIST as assessed by RNA-FISH (53) and respond to retinoic acid (RA)-mediated differentiation cues to give rise to neuronal progenitors (54). Therefore, this cell line provides a good system to probe for the dynamic pattern of *XIST* promoter activity during the initial expression of XIST RNA. HEK293T is an epithelial cell line of female origin, exhibits consistent expression of XIST denoting the maintenance phase of XCI (55).

XIST expression is detected only in the female cell line HEK293T and not in the cell lines of male origin - NT2/D1 cells and DLD1 (epithelial cell line) **(Figure 1A)**. To understand the regulation of *XIST* during the initiation phase, we performed RA-mediated differentiation of NT2/D1 cells and show that the levels of pluripotency factors – OCT4 (POU5F1), SOX2, NANOG decline, and the expression of neuronal progenitor specific marker – PAX6 is upregulated during differentiation, thus corroborating the previous report (54) **(Figure 1B,C)**. In this differentiation paradigm of NT2/D1 cells, XIST expression is upregulated progressively **(Figure 1D, E, F)**. RNA FISH analysis indicated that over 20% NT2/D1 cells exhibit robust XIST RNA FISH signal upon differentiation of NT2/D1 for 5 days **(Figure 1E, F)**. We verified that NT2/D1 cells harbour two intact and one broken X chromosome copies by performing X chromosome paint for metaphase spreads as well as interphase nuclei **(Figure 1E)**. It is important to note that the pattern remains the same in undifferentiated and 5-day RA-treated cells. Despite harbouring two intact X chromosomes, only about 20% of NT2/D1 cells differentiated for 5 days show XIST RNA FISH signal. Whether this is due to differential kinetics of XIST RNA expression in NT2/D1 or non-persistent expression and/or maintenance of XIST RNA as seen in various hESC lines (56–59) awaits further investigation. Nevertheless, our aim was to understand the transcriptional regulation of XIST in the context of initiation phase of XCI and based on the results presented in **Figure 1A-F**, we believe that NT2/D1 cells can serve as a good model to address this question, especially since hESCs have not served as an ideal model for the purpose so far.

**Figure 1.**
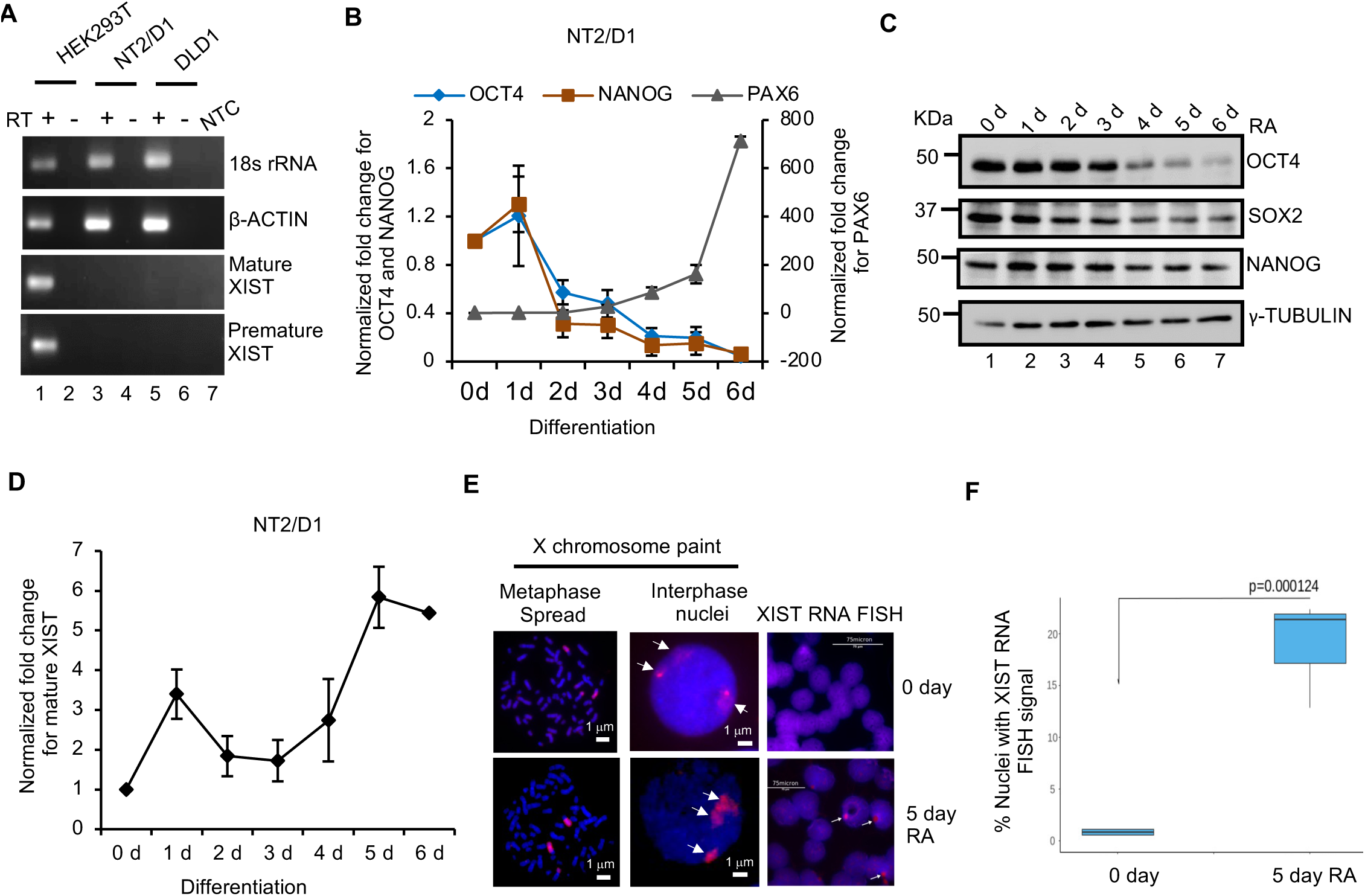
NT2/D1 and HEK293T cells provide the contexts for initiation and maintenance phases of XCI. **(A)** Semi-quantitative RT-PCR for mature and premature XIST using cDNA prepared from HEK293T (female), NT2/D1 (male) and DLD1 (male) cells. 18s rRNA and β-ACTIN serve as controls. **(B)** qRT-PCR depicting a decrease in the levels of OCT4 and NANOG and increase in PAX6 levels upon RA-mediated differentiation of NT2/D1 cells for 6 days. X-axis represents the differentiation time-point and Y-axis represents the fold change normalized to 18s rRNA. Each point on the graph represents values from 5 independent experiments and error bar represents ±S.E.M. **(C)** Immunoblotting showing a decrease in OCT4, SOX2 and NANOG levels upon RA-mediated differentiation of NT2/D1 cells for 6 days. γ-TUBULIN serves as an equal loading control. **(D)** qRT-PCR depicting an increase in the levels of XIST upon RA-mediated differentiation of NT2/D1 cells for 6 days. X-axis represents the differentiation time-point and Y-axis represents the fold change normalized to 18s rRNA. Each point on the graph represents values from 3 independent experiments and error bar represents ±S.E.M. **(E)** X chromosome paint for metaphase spread and interphase nuclei from undifferentiated (0 day) and 5day RA-treated NT2/D1cells. RNA FISH for mature *XIST* for undifferentiated (0 day) and differentiating (5day, RA) NT2/D1 cells. Arrowheads indicate the FISH signal. **(F)** Quantification for the RNA FISH signals in 0 day and 5day RA treated NT2/D1 cells. N=3, 200 nuclei were counted for each replicate and statistical significance was ascertained by Student’s T-test.

### Transcription from *XIST* promoter is governed by the promoter as well as exon 1 of *XIST*

The pioneering study attempting to characterize the promoter of human *XIST* restricted their analysis to +50 to −50 bp from the TSS, since it was found to be conserved across four mammalian species (50). We chose to test the larger promoter region to uncover the unique potential regulatory elements for human *XIST* by cloning upto 4408 bp upstream of *XIST* TSS into a promoter-less luciferase reporter vector **(Figure 2A)**. These DNA constructs were then transfected into NT2/D1 and HEK293T cells and assayed for the presence of promoter by measuring the induced firefly luciferase activity. Similar to the first study on characterizing *XIST* promoter elements (50), we observed that the +50 bp to −51 bp region was sufficient to drive the transcription of luciferase gene **(Figure 2B,C)**. Also, the promoter activities for all other fragments with increasing distance from the TSS (except for the fragment +50 bp to −260 bp, bar 3 in **Figure 2B,C**) remained constant when compared to the vector control. As a control, +50 bp to −1050 bp was cloned in the anti-sense (AS) orientation. Since *XIST* promoter is unidirectional, the fragment cloned in the AS direction failed to transcribe the reporter gene and hence did not exhibit any measurable reporter activity (AS, **Figure 2B,C)**. We also measured the reporter activities in differentiating NT2/D1 cells (0 to 2 days) and observed a significant decrease in luciferase levels **(Figure 2D)**. Expression of previously identified transcriptional activators of *XIST* – SP1 and YY1 (50) also decreased upon differentiation of NT2/D1 cells, reaffirming their roles in regulating XIST **(Figure 2E)**.

**Figure 2.**
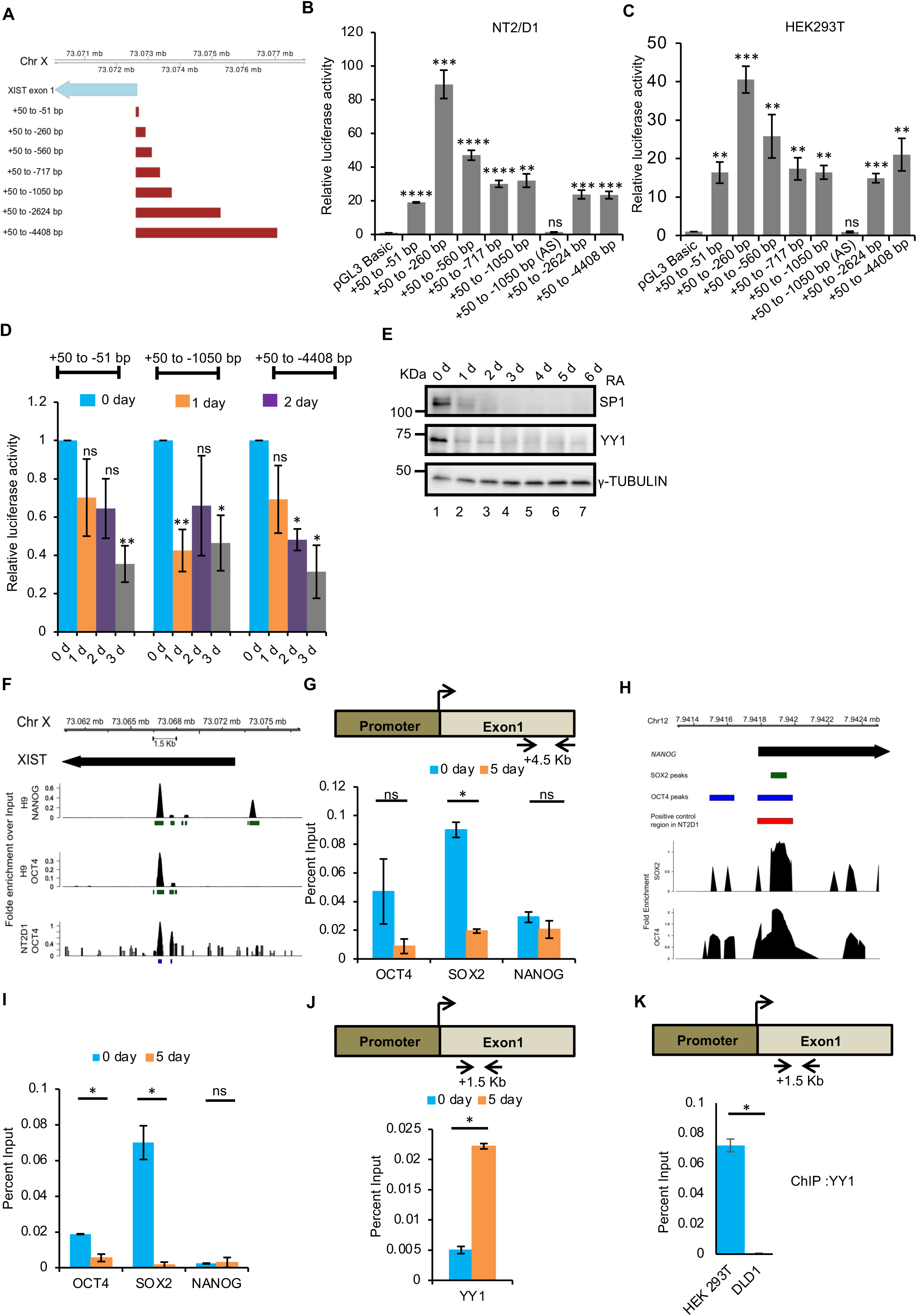
Induction of XIST during differentiation of NT2/D1 cells is governed by the promoter as well as exon1 of *XIST*. **(A)** The brown boxes denote the fragments of the region upstream to XIST clone into pGL3 Basic Luciferase reporter for testing potential promoter activity. The blue arrow indicates the first exon of *XIST* with the arrowhead denoting the direction of transcription and the tail indicating the TSS. The numbers flanking each brown box denote the relative location of the fragment with respect to the Transcription Start Site (TSS) of XIST **(B,C)** Luciferase reporter activities for *XIST* promoter constructs transfected into NT2/D1 cells (B) or HEK293T (C) compared to pGL3 Basic. Firefly luciferase activities represented here are normalized to Renilla luciferase activity, which serves as an internal control. X-axis indicates DNA constructs transfected and Y-axis represents the normalized fold change in firefly luciferase activity. Each bar represents values from three independent experiments. Error bar represent ±S.E.M. Asterisks represent the significance over vector control as per Student’s T-test (****P value < 0.0001, ***P value < 0.001, **P value < 0.01, *P value < 0.05, ns = non-significant). **(D)** Luciferase reporter activities of +50 bp to −51 bp, +50 bp to −1050 bp and +50 bp to −4408 bp promoter constructs decrease upon RA-mediated differentiation of NT2/D1 for 3 days. Firefly luciferase activities represented here are normalized to Renilla luciferase activity, which serves as an internal control. X-axis indicates differentiation time-points and Y-axis represents the normalized fold change in the firefly luciferase activity. Each bar represents values from three independent experiments. Error bar represents ±S.E.M. Asterisks represent the significance over vector control as per Student’s T-test (**P value < 0.01, *P value < 0.05, ns = non-significant). **(E)** SP1 and YY1 proteins decrease upon RA-mediated differentiation of NT2/D1 for 6 days. γ-TUBULIN serves as a loading control. **(F)** ChIP-seq peaks for OCT4 (SRR5642847) and NANOG (SRR5642845) for the *XIST* locus in human embryonic stem cell line H9 and OCT4 ChIP-seq peak for NT2/D1 cells (SRR1640282). The arrowhead indicates direction of *XIST* transcription. All peaks were confirmed to have a p value < 0.05 as reported by MACS2 callpeak function **(G)** ChIP-qPCR analysis showing a decrease in OCT4, SOX2 and NANOG enrichment on the *XIST* at +4.5 Kb (as shown in the schematic above) in undifferentiated (0 day) versus 5 day differentiated NT2/D1 cells. **(H)** Positive control for the ChIP of OCT4/POU5F1 in NT2D1. The positive control is selected based on the location of two existing peaks in NT2D1 cells. **(I)** ChIP-qPCR analysis for the control region showing enrichment of pluripotency factors on NANOG promoter. Each bar represents values from 2 independent experiments. Error bar represent the ±S.E.M. Asterisks represent the significance over DLD1 as per Student’s T-test (*P value < 0.05, ns=non-significant) **(J)** ChIP-qPCR analysis demonstrating a change in the occupancies YY1 on *XIST* promoter-proximal region (+1.5 Kb) (as shown in the schematic above) in undifferentiated (0 day) versus 5 day differentiated NT2/D1 cells. X-axis represents the immunoprecipitated factor and Y-axis represents the enrichment calculated as percent input. Each bar represents values from 2 independent experiments. Error bar represents ±S.E.M. Asterisks represent the significance over undifferentiated cells (0 day) as per Student’s T-test (*P value < 0.05, ns = non-significant). **(K)** ChIP-qPCR analysis for YY1 on *XIST* promoter proximal region (∼ +1.5 Kb) (as shown in the schematic above) in HEK293T (female, blue bar) and DLD1 (male, brown bar) cells. Each bar represents values from 3 independent experiments. Error bar represents ±S.E.M. Asterisks represent the significance as per Student’s T-test (*P value < 0.05).

The tested promoter fragments exhibited a similar trend in both the cell lines tested. This implies that the assessed transcription activity is independent of the cellular context wherein all the necessary transcription factors are present in both cell types. This raises the question as to why *XIST* fails to express in the undifferentiated NT2/D1 cells. In mouse system, the pluripotency factors negatively regulate *Xist* and therefore NT2/D1 cells, which mimic the undifferentiated embryonic stem cells and exhibit a negative correlation between *XIST* and pluripotency factor expression, were the logical candidates for assessing their role. Towards this, we scanned the *XIST* locus for binding sites of pluripotency factors by re-analysing the available ChIP-sequencing datasets for OCT4 and NANOG in H9 hESC and OCT4 in NT2/D1 cells and observed their enrichment on the 1^st^ exon of *XIST*, ∼ +4.5 Kb from the TSS in both the cell types independent of sex **(Figure 2F)**. We validated their binding at this specific location by ChIP-qPCR and observed an enrichment of OCT4 and significant enrichment of SOX2 in undifferentiated (0 day) as compared to 5 day differentiated NT2/D1 cells **(Figure 2G)**. This indicates that binding of the pluripotency factors might serve to repress *XIST* in an undifferentiated state. *NANOG* promoter region comprising ChIP-seq peaks for OCT4 and SOX2 served as a positive control for pluripotency factor ChIP-qPCR assay **(Figure 2H,I)**.

The next prominent question was to identify the factors positively regulating *XIST* in this scenario. YY1 has been shown to bind at +1.5 Kb site from the TSS and serve as a transcriptional activator of XIST (51, 52). Therefore, we determined YY1 binding at this site by ChIP-qPCR and observed a significant enrichment for day 5 differentiated NT2/D1 cells **(Figure 2J)**. We also confirmed YY1 enrichment at this site specifically in the female cells by comparing ChIP-qPCR between female cells (HEK293T) and male cells (DLD1) **(Figure 2K)**. These results suggest that the enrichment of the pluripotency factors 4.5 Kb downstream of the TSS negatively correlates with XIST expression. Occupancy of YY1, the known transcriptional activator of *XIST* at +1.5 Kb correlates with the induction of *XIST* in NT2/D1 differentiation model. Therefore, it seems likely that similar to the mouse system, the pluripotency factors act as the potential repressors of human *XIST*.

### Pluripotency factors negatively regulate *XIST*

To validate if pluripotency factors indeed repress *XIST*, two approaches were employed – (1) Depleting their levels in NT2/D1 cells, which express high levels of OCT4, SOX2, NANOG, and do not express XIST and hence providing the initiation phase context, (2) Forced overexpression of pluripotency factors in HEK293T cells that do not endogenously express these factors and exhibit high expression of XIST (maintenance phase). Knockdown of OCT4, SOX2 or NANOG in undifferentiated NT2/D1 cells **(Figure 3A)** led to a significant increase in XIST levels **(Figure 3C)** when either OCT4 alone was depleted or both SOX2 and NANOG were depleted **(Figure 3C)**. Increase in the expression of the neuronal marker – PAX6 is expected upon knockdown of pluripotency factors and hence serves as a control **(Figure 3B)**. We would like to point out that a significant upregulation in XIST expression was observed only when the levels of all the three factors were highly reduced, which is under the condition where OCT4 is knocked down.

**Figure 3.**
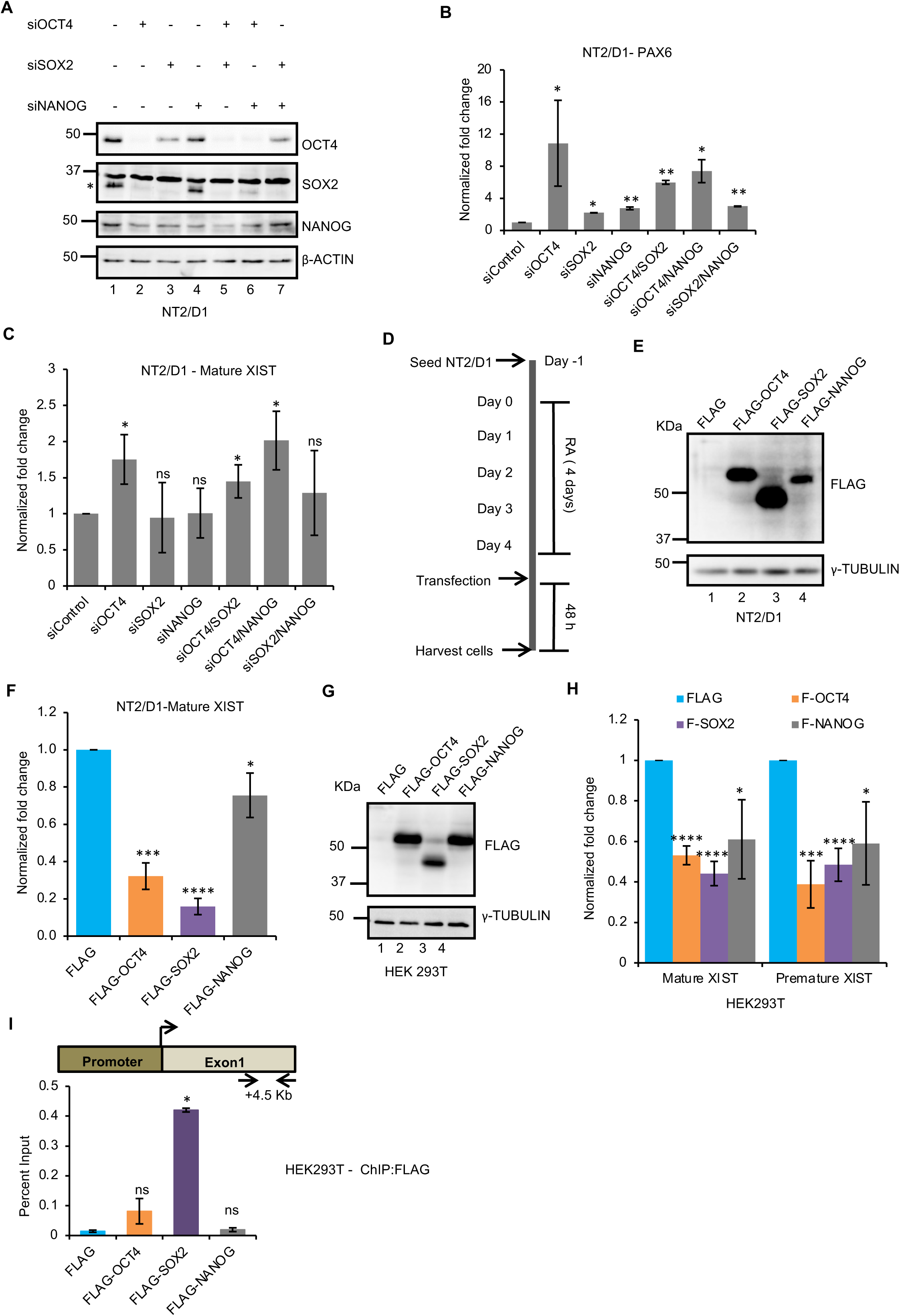
Pluripotency factors repress *XIST* by binding to exon 1 (+4.5 Kb) site. **(A)** Immunoblotting to determine the knockdown efficiencies of OCT4, SOX2 and NANOG in NT2/D1 cells. β-ACTIN serves as an equal loading control. **(B)** qRT-PCR for PAX6 upon siRNA mediated knockdown of OCT4, SOX2, NANOG in NT2/D1 cells. X-axis represents siRNA transfected and Y-axis represents the fold change normalized to 18s rRNA. Each bar represents values from 3 independent experiments. Error bar represents the ±S.E.M. Asterisks represent the significance over vector control as per Student’s T-test (**P value < 0.01, *P value < 0.05, ns=non-significant). **(C)** qRT-PCR for mature XIST upon siRNA mediated knockdown of OCT4, SOX2, NANOG in NT2/D1 cells. X-axis represents siRNA transfected and Y-axis represents the fold change normalized to 18s rRNA. Each bar represents values from 3 independent experiments. Error bar represents the ±S.E.M. Asterisks represent the significance over vector control as per Student’s T-test (**P value < 0.01, *P value < 0.05, ns=non-significant). **(D)** Experimental scheme to overexpress OCT4, SOX2, NANOG in NT2/D1 cells differentiated for 4 days. **(E)** Immunoblotting for FLAG to confirm the over-expression of OCT4, SOX2, NANOG in NT2/D1 cells differentiated for 4 days. γ-TUBULIN serves as an equal loading control. **(F)** qRT-PCR showing a significant reduction in mature XIST upon over-expression of pluripotency factors in NT2/D1 cells differentiated for 4 days. X-axis represents transfected DNA and Y-axis represents the fold change normalized to 18s rRNA. Each point on the graph represents values from 3 independent experiments and error bar represents ±S.E.M. Asterisks represent the significance over vector control as per Student’s T-test (****P value < 0.0001, P value < 0.001, P value < 0.05). **(G)** Immunoblotting for FLAG to confirm the over-expression of OCT4, SOX2, NANOG in HEK293T cells. γ-TUBULIN serves as an equal loading control. **(H)** qRT-PCR showing a significant reduction in mature and premature XIST upon over-expression of pluripotency factors in HEK293T cells. X-axis represents the mature or premature XIST and Y-axis represents the fold change normalized to 18s rRNA. Each point on the graph represents values from 5 independent experiments and error bar represents ±S.E.M. Asterisks represent the significance over vector control as per Student’s T-test (****P value < 0.0001, ***P value < 0.001, *P value < 0.05). **(I)** ChIP-qPCR showing occupancies of OCT4, SOX2, NANOG on the exon 1 (+4.5 Kb) site upon their over-expression in HEK293T cells. X-axis represents the transfected DNA and Y-axis represents the enrichment calculated as percent input. Each point on the graph represents values from 2 independent experiments and error bar represents ±S.E.M. Asterisks represent the significance over vector control as per Student’s T-test (*P value < 0.05).

To further test the repressive role of pluripotency factors, we overexpressed OCT4, SOX2 or NANOG in 4-day differentiated NT2/D1 cells as per the scheme depicted in **Figure 3D**. 4-day differentiation time point was chosen since pluripotency factors begin to decline **(Figure 1 B,C)** and XIST expression is induced **(Figure 1D)** at this time point. A significant decline in XIST expression was observed under these experimental conditions, thereby strengthening the proposed repressive role for OCT4, SOX2 and NANOG **(Figure 3E,F)**. These results led us to conclude that OCT4, SOX2 and NANOG act as the repressors of *XIST* in embryonic stem cells, wherein it is necessary to prevent premature induction of XIST.

The counter experiment of ectopically overexpressing OCT4 or SOX2 or NANOG in HEK293T **(Figure 3G)**, led to a significant decrease in mature XIST expression assessed by qRT-PCR primers designed at exon-exon junction **(Figure 3H)**. A similar trend of decrease was obtained for premature XIST, assessed by primers targeting intronic region, which suggests that the effect observed is indeed a transcriptional and not a post-transcriptional effect **(Figure 3H).** The overexpressed OCT4 and SOX2 proteins are enriched at the +4.5 Kb region on the exon 1 of *XIST*, providing a clear evidence that the pluripotency factors negatively regulate transcription of *XIST* by directly binding to this site **(Figure 3I)**. Considering the effect of OCT4, SOX2 and NANOG on XIST expression in HEK293T cells, we postulate that occupancy of the +4.5 Kb site by the pluripotency factors could exert a more potent effect on the expression of XIST in conjunction with the binding of the known activator YY1 on the promoter-proximal region.

### The pluripotency factor bound regulatory site on *XIST* exon 1 is a potential *cis* regulatory element (cRE)

The results obtained thus far provided compelling evidence to further characterize the regulatory region of the human *XIST* exon 1 (+4.5 Kb from the TSS). It seemed likely that this element harbours regulatory potential whose function is modulated by the binding of pluripotency factors. To assess the functional significance of this site, we performed ChIP-qPCR for active (H3K27ac) and inactive (H3K27me3) histone modifications in female (HEK293T) as well as male (DLD1) cells and observed a significant enrichment of active chromatin marks only in the female cells indicating that we are indeed scoring for the *XIST* locus from the Xi **(Figure 4A)**. Additionally, there was no significant difference between H3K27me3 enrichment on this site in the male and female cells suggesting it to be enriched on the *XIST* locus of Xa **(Figure 4A)**. Another feature of the newly identified cRE is the significant enrichment of CTCF specifically in female cells (MCF7, HeLa, HEK 293 and female skin epithelium) versus male (DLD1 and male skin epithelium) cells as evident from the analysis of available ChIP-sequencing datasets (60, 61) **(Figure 4B)**. The inclusion of primary cell data lends confidence in the findings and excludes the possibility of observing CTCF occupancy merely as an artefact of cultured cell lines. A previous report ruled out the role of CTCF in regulating *XIST* since they observed no appreciable difference in its enrichment on the part of *XIST* locus tested between male and female cells (51). Nonetheless, this report has not examined the site identified in our study and hence, we investigated its role in governing *XIST* transcription.

**Figure 4.**
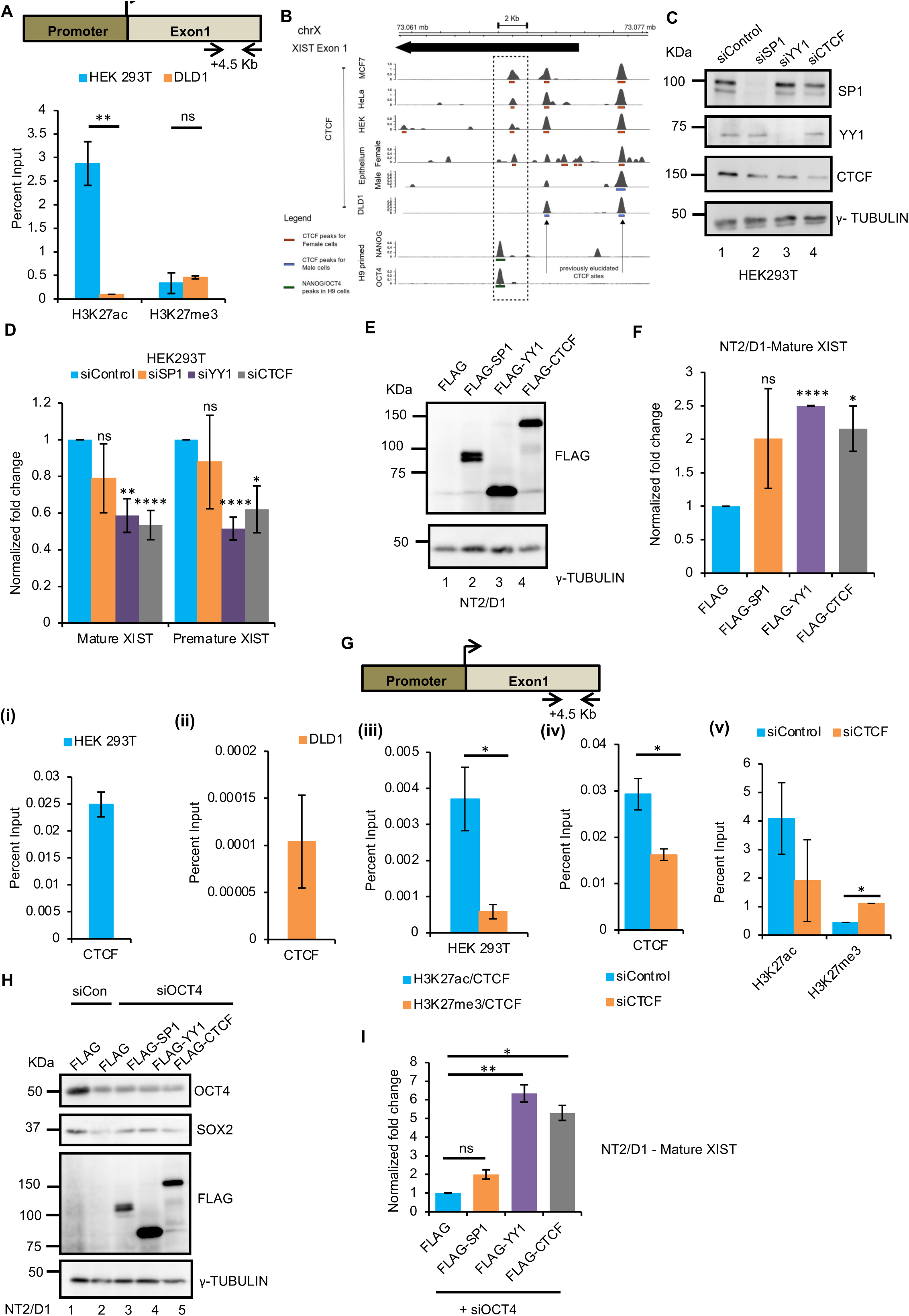
Pluripotency factor binding element is the potential cRE. **(A)** ChIP-qPCR analysis showing enrichment of active histone mark H3K27ac and repressive histone mark H3K27me3 for *XIST* cRE (+4.5 Kb, as shown in the schematic above) on exon1 in HEK293T (female, blue bar) and DLD1 (male, brown bar) cells. X-axis represents the antibodies used for ChIP and Y-axis represents the enrichment calculated as percent input. Each bar represents values from 3 independent experiments. Error bar represent ±S.E.M. Asterisks represent the significance over DLD1 ChIP as per Student’s T-test (*P value < 0.05, ns = non-significant). **(B)** ChIP-seq peaks for OCT4 (SRR5642847) and NANOG (SRR5642845) for the *XIST* locus in human embryonic stem cell line H9 and OCT4 ChIP-seq peak for NT2/D1 cells (SRR1640282). CTCF ChIP-seq for male cell lines - DLD1 (DRR014660), skin epithelium (SRR6213724) and female cell lines - MCF7 (SRR577680, SRR577679), HeLa (SRR227659, SRR227660), HEK293 (DRR014670), skin epithelium (SRR6213076) lines. The arrowhead indicates the direction of *XIST* transcription and tail denoted the TSS. All peaks were confirmed to have a p value < 0.05 as reported by MACS2 callpeak function. **(C)** Immunoblotting to confirm siRNA mediated knockdown of SP1, YY1, CTCF in HEK293T cells. γ-TUBULIN serves as an equal loading control. **(D)** qRT-PCR demonstrating reduction in XIST (both mature and premature) levels upon knockdown of YY1 or CTCF in HEK293T cells. X-axis represents the mature or premature XIST, and Y-axis represents the fold change normalized to 18s rRNA. Each point on the graph represents values from 4 independent experiments and error bar represents ±S.E.M. Asterisks represent the significance over vector control as per Student’s T-test (****P value < 0.0001, **P value < 0.01, *P value < 0.05, ns=non-significant). **(E)** Immunoblotting to confirm over-expression of SP1, YY1, CTCF in NT2/D1 cells. γ-TUBULIN serves as an equal loading control. **(F)** qRT-PCR demonstrating a significant increase in XIST expression upon over-expression of YY1 or CTCF in NT2/D1 cells. X-axis represents the transfected DNA and Y-axis represents the fold change normalized to 18s rRNA. Each point on the graph represents values from 3 independent experiments and error bar represents ±S.E.M. Asterisks represent the significance over vector control as per Student’s T-test (****P value < 0.0001, *P value < 0.05, ns=non-significant). **(G)** ChIP-qPCR analysis showing enrichment of CTCF on *XIST* cRE (+4.5 Kb) on exon1 in (i) HEK293T (female, blue bar) and (ii) DLD1 (male, brown bar) cells (iii) Sequential ChIP in HEK293T cells (iv-v) ChIP-qPCR analysis for CTCF (iv) and H3K27ac, H3K27me3 (v) upon siRNA mediated knockdown of CTCF in HEK293T cells (brown bars). Each bar represents values from 3 (for (i)) or 2 (for (ii-v)) independent experiments. Error bar represent the ±S.E.M. Asterisks represent the significance as per Student’s T-test (*P value < 0.05). **(H)** Immunoblotting for OCT4 and SOX2 upon the knockdown of OCT4 and for FLAG to confirm the over-expression of SP1, YY1, CTCF in NT2D1 cells. γ -TUBULIN serves as a loading control. **(I)** qRT-PCR for mature XIST upon knockdown of OCT4 and over-expression of SP1, YY1, CTCF in NT2D1 cells. X-axis represents DNA and siRNA transfected and Y-axis represent the fold change normalized to 18s rRNA. Each bar represents values from 2 independent experiments. Error bar represent the ±S.E.M. Asterisks represent the significance over vector control as per Student’s T-test (**P value < 0.01, *P value < 0.05, ns = non-significant).

Perturbing the levels of SP1, YY1, and CTCF in HEK293T cells caused a significant decrease in both mature and premature XIST levels upon knockdown of YY1 or CTCF **(Figure 4C,D)**. Overexpression of SP1, YY1 or CTCF in undifferentiated NT2/D1 cells which expresses XIST at very low levels led to a significant increase in XIST levels upon over-expression of YY1 or CTCF **(Figure 4E,F)**. These results strongly suggest that in addition to YY1, CTCF is an essential regulator of *XIST*. Furthermore, we validated CTCF ChIP-sequencing analysis finding by performing ChIP-qPCR for CTCF in HEK293Tand DLD1 cells **(Figure 4G (i,ii))** and observed enrichment on the cRE in the female cells by an order of magnitude which positively correlates with XIST expression **(Figure 4D)**. To confirm whether CTCF is indeed bound to cRE only on the Xi, we performed sequential ChIP-qPCR in HEK293T cells using anti-H3K27ac antibody (mark enriched at +4.5 Kb site only in female cells as per Fig. 4A) or anti-H3K27me3 antibody (mark enriched at +4.5 Kb site in both female and male cells as per Fig. 4A) antibodies for the first ChIP followed by second ChIP using anti-CTCF antibody. We observed a higher enrichment of CTCF when the first ChIP was performed using H3K27ac antibody **(Figure 4G (iii)**, **blue bar)** compared to H3K27me3 antibody **(Figure 4G (iii), brown bar)**. Further confirmation for the direct role of CTCF in regulating XIST transcription is evident by the decreased enrichment of CTCF **(Figure 4G (iv))** and increased H3K27me3 levels at the cRE observed upon knockdown of CTCF in HEK293T cells **(Figure 4G (v))**.

We investigated the interplay between transcriptional activators and repressors of XIST by overexpressing potential transcriptional activators - SP1 or YY1 or CTCF coupled with the knockdown of OCT4 in NT2/D1 cells. Interestingly, we observed a significantly higher upregulation of XIST when the overexpression of YY1 or CTCF was coupled with the knockdown of OCT4 **(Figure 4H,I)**, when compared with only the over-expressions of YY1 or CTCF **(Figure 4E,F)** or the knockdown of pluripotency factors alone **(Figure 3A,B)**. This suggests that the balance of activators (YY1, CTCF) and repressors (OCT4, SOX2, NANOG) govern the transcriptional activation of *XIST*. It is noteworthy that the siRNA-mediated knockdown of OCT4 depletes not only OCT4 but also SOX2 **(Figure 4H)**. Altogether, these results indicate the existence of a cRE on exon 1 of *XIST* whose function is modulated by CTCF or pluripotency factors, which in turn determines transcriptional status of *XIST*

### CTCF binding to cRE assists in stabilizing YY1 binding at the *XIST* promoter-proximal site

YY1 has been demonstrated to be a conserved transcriptional activator of XIST (51, 52), which is also recapitulated in our study. In order to test the functional significance of CTCF in regulating *XIST*, we assessed occupancy of YY1 at previously identified promoter-proximal region (+1.5 Kb) upon knockdown of CTCF. Interestingly, while the levels of YY1 do not change upon depleting CTCF in HEK293T cells **(Figure 4C)**, a significant decrease in its enrichment can be observed at the promoter-proximal region **(Figure 5A)**. Furthermore, we show that the downregulation in XIST observed upon depleting CTCF can be rescued by overexpressing YY1 in HEK293T cells **(Figure 5B,C)**. This suggests that CTCF assists either in the recruitment or maintenance of YY1 binding at the promoter-proximal region. Based on these results, we envisaged the significance of CTCF binding to cRE as an important determinant of XIST transcription. To test this possibility, we employed the CRISPR-dCas9-KRAB directed approach to interfere with CTCF binding at the cRE **(Figure 5D**) and observed a significant downregulation in mature as well as premature XIST levels **(Figure 5E)**. ChIP-qPCR analysis indicate enrichment of dCas9 (monitored by FLAG ChIP) and H3K9me3 mark at cRE suggestive of efficient targeting **(Figure 5F)**. This led to a concomitant decrease in CTCF occupancy at cRE **(Figure 5F)**. In concordance with our CTCF knockdown results **(Figure 5A)**, YY1 occupancy at the promoter-proximal region declined under these conditions as well **(Figure 5G)**. We also scored for H3K9me3 at the promoter-proximal region to determine if the observed XIST downregulation upon targeting CRISPR-dCas9-KRAB to cRE is an outcome of spreading of repressive mark to the promoter. Surprisingly, we observed a decrease in H3K9me3 mark at the promoter-proximal region. The reason for this is not fully understood and needs further investigation. Based on these results, we infer that binding of CTCF to the newly identified cRE regulates *XIST* in a YY1-dependent manner.

**Figure 5.**
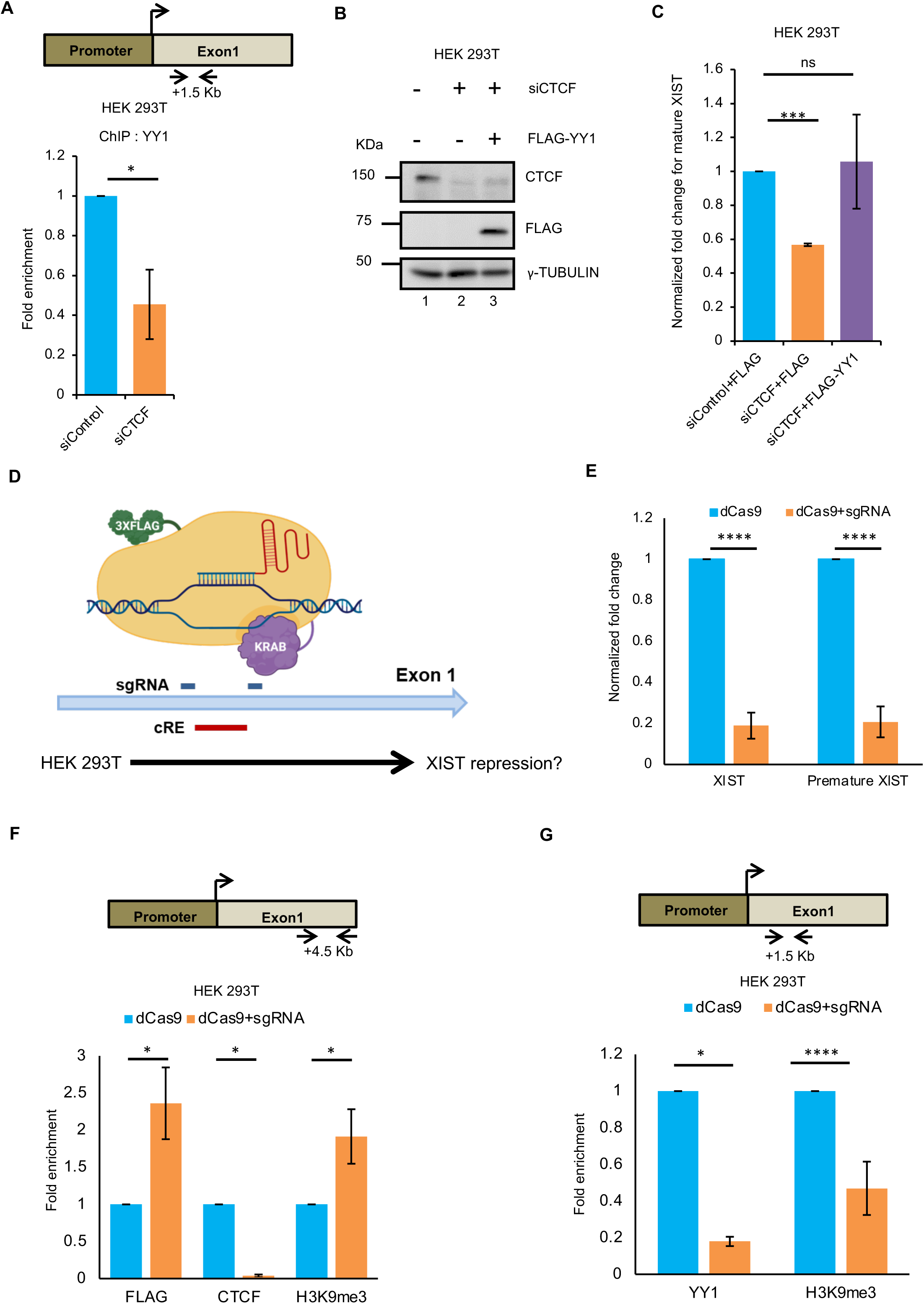
CTCF assists in the recruitment or maintenance of YY1 binding to the promoter-proximal region of *XIST*. **(A)** ChIP-qPCR analysis showing a reduction of YY1 occupancy on the promoter-proximal site (+1.5 Kb) upon siRNA mediated knockdown of CTCF in HEK293T cells (brown bar). X-axis represents the antibody used for ChIP, and Y-axis represents the normalized fold enrichment over control siRNA (blue bar). Each bar represents values from 3 independent experiments. Error bar represents ±S.E.M. Asterisks represent the significance over YY1 ChIP for vector control as per Student’s T-test (*P value < 0.05). X-axis represents the transfected siRNA, and Y-axis represents the normalized fold enrichment over control siRNA. **(B)** Immunoblotting to confirm knockdown of CTCF and over-expression of FLAG-tagged YY1 in HEK293T cells. γ-TUBULIN serves as an equal loading control. **(C)** qRT-PCR for mature XIST upon knockdown of CTCF and/or over-expression of FLAG-YY1 in HEK293T cells. X-axis represents the DNA and siRNA transfected and Y-axis represents the normalized fold enrichment over 18s rRNA. Each bar represents values from 2 independent experiments. Error bar represents ±S.E.M. Asterisks represent the significance over control (1^st^ bar) as per Student’s T-test (***P value < 0.001, ns=non-significant). **(D)** Schematic depicting dCas9-KRAB based repression strategy. **(E)** qRT-PCR demonstrating reduction in XIST (both mature and premature) levels upon transfecting dCas9+sgRNA in HEK293T cells. X-axis represents the mature or premature XIST, and Y-axis represents the fold change normalized to dCas9 only control. Each point on the graph represents values from 6 independent experiments and error bar represents ±S.E.M. Asterisks represent the significance over vector control as per Student’s T-test (****P value < 0.0001) **(F)** ChIP-qPCR analysis showing enrichment of dCas9, CTCF and H3K9me3 at cRE (+4.5 Kb, as shown in the schematic above) on exon1 in HEK293T cells transfected with just dCas9 (blue bar) or dCas9+sgRNA (brown bar). X-axis represents the antibodies used for ChIP and Y-axis represents the enrichment calculated as percent input. Each bar represents values from 3 independent experiments. Error bar represent ±S.E.M. Asterisks represent the significance over dCas9 control as per Student’s T-test (*P value < 0.05). **(G)** ChIP-qPCR analysis showing enrichment of YY1 and H3K9me3 at promoter-proximal region (+1.5 Kb, as shown in the schematic above) on exon1 in HEK293T cells transfected with just dCas9 (blue bar) or dCas9+sgRNA (brown bar). X-axis represents the antibodies used for ChIP and Y-axis represents the enrichment calculated as percent input. Each bar represents values from 3 independent experiments. Error bar represent ±S.E.M. Asterisks represent the significance over dCas9 control as per Student’s T-test (****P value < 0.0001, *

### Discussion

*XIST* is the critical regulator of XCI, a mechanism to correct for the gene dosage imbalance between males and females of the placental mammals (1–4). Being central to the process of XCI, it is essential to modulate XIST levels temporally to ensure faithful execution of the process and hence the developmental programs. Despite significant conservation of X-linked genes, chromosomal synteny as well as the process of XCI guided by XIST lncRNA between several eutherian mammals, XCI is initiated in diverse ways in different species. One of the possible explanations could be evolutionary alterations in the developmental programs across species necessitating robustness in the regulation of XCI to accommodate these changes (46). Extensive studies using the mouse system have highlighted the multiple ways in which the transcription of *Xist* is regulated during the initiation and the maintenance phases of XCI. Thus far, only YY1 has been discovered to be the common factor regulating *XIST* in mouse and human (51, 52). Here, we have uncovered a complex regulatory network involving pluripotency factors, YY1 and CTCF that shapes the transcriptional outcome from the *XIST* promoter **(Figure 6)**.

**Figure 6.**
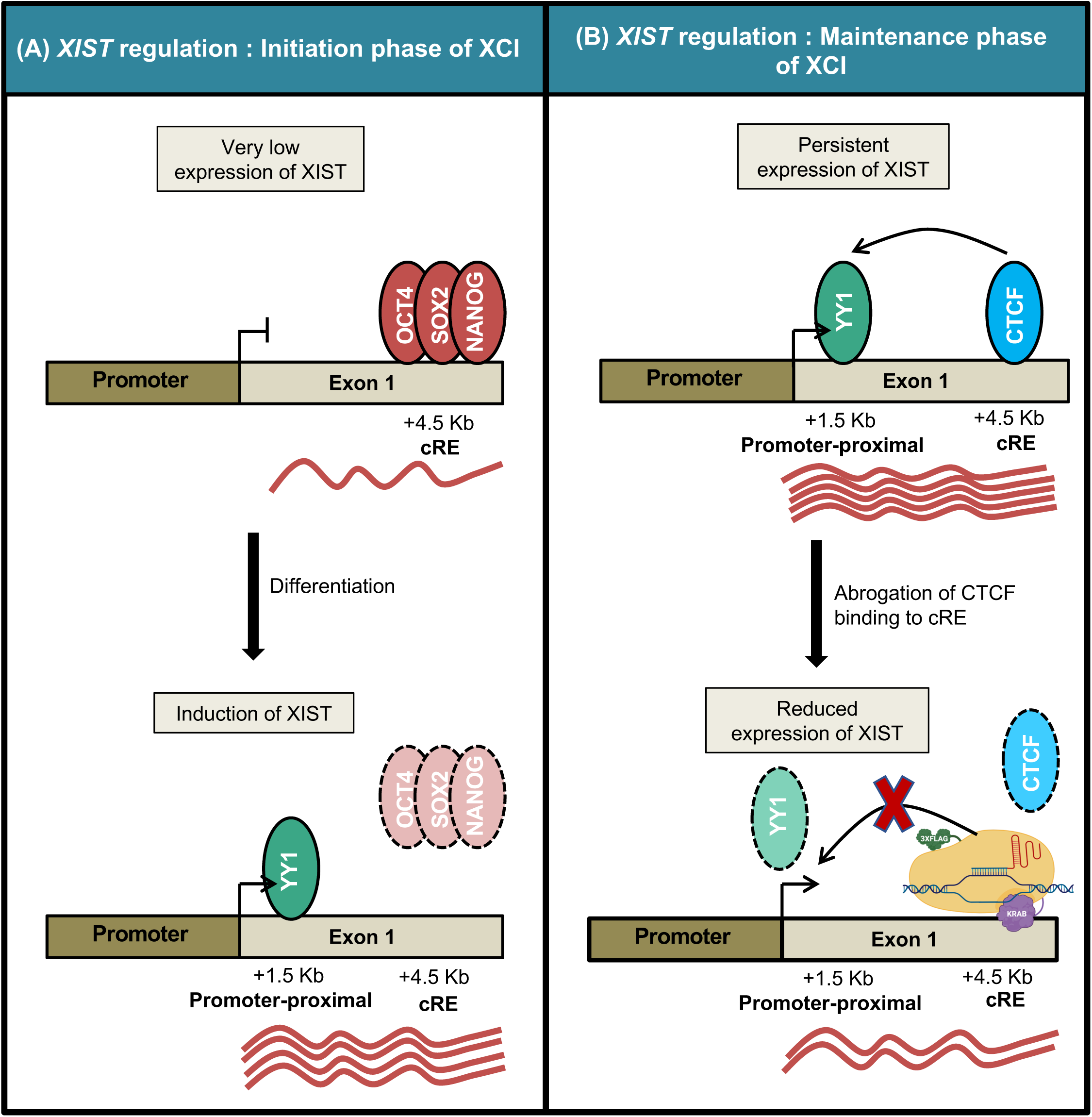
A model illustrating the role of CTCF bound cRE in dictating the transcription from *XIST* promoter. (**A**) In undifferentiated ES cells, cRE (+4.5 Kb) of *XIST* is bound by the pluripotency factors (OCT4, SOX2, NANOG) keeping XIST repressed. Upon differentiation, levels as and enrichment of pluripotency factors on the cRE decreases. Subsequently, YY1 now occupies the promoter-proximal site leading to induction of XIST. (**B**) In differentiated cells, CTCF binding to cRE (+4.5 Kb) enables either the recruitment or maintenance of YY1 at the promoter proximal region, maintaining persistent expression of XIST. Abrogation of CTCF binding to cRE by CRISPR-dCas9-KRAB interference disrupts YY1 binding to the promoter-proximal region causing downregulation of XIST.

Unlike mouse ES cells, the human counterpart has not served as a tractable model to address the aspects of XCI and *XIST* regulation. Multiple groups have reported considerable variation in XIST expression, as well as XCI status between human ES lines over passages (57, 62). However, some recent studies describe a method of culturing human ES cell lines, exhibiting consistent XIST expression as well as XCI upon differentiation (63, 64). The difficulty of human ES cells as a model notwithstanding, we have worked with NT2/D1 and HEK293T as models of initiation and maintenance phases of XCI respectively. Moreover, we have reanalysed the data relevant to our study from multiple cell types including a human ES line and primary tissues of the female and male origin to discover physiologically relevant Xi-specific factors contributing to *XIST* regulation.

To address *XIST* regulation in the context of initiation phase of XCI, we have extensively worked with NT2/D1 cells, a human male embryonic carcinoma cell line. The NT2/D1 cell line provides the context for the initiation phase of XCI since it harbours properties similar to the embryonic stem cells and has been extensively used to understand the human stem cell biology (65–67). More importantly, it also expresses low levels of XIST (53), analogous to that seen for human and mouse ES cells as well blastocyst (5, 46, 47, 68–70). We show for the first time that XIST expression can be induced in these cells upon RA-mediated differentiation and hence, NT2/D1 can serve as a model to determine the induction of *XIST*. The fact that there is a negative correlation between *XIST* induction and loss of OCT4, SOX2 and NANOG during differentiation, prompted us to speculate their role in governing XIST transcription. Moreover, there have been studies reporting pluripotency factors to be important positive and negative regulators of mouse *Xist* and *Tsix* respectively (33, 35, 36). Upon reanalysing published ChIP-sequencing datasets and the experimental validation, we provide evidence for the first time that pluripotency factors repress *XIST*. Interestingly, although levels of YY1, a known transcriptional activator of *XIST*, decline during differentiation, its occupancy on the previously reported promoter-proximal site (+1.5 Kb) increases, causing a significant increment in XIST expression by day 5 of differentiation (51, 52). We believe that regulation mediated by the pluripotency network could be an essential way to control upregulation of XIST temporally. These findings provide compelling evidence suggestive of this gene body regulatory region to be a potential Xi-specific *cis* regulatory element. Indeed, a higher enrichment of activation associated histone marks – H3K27ac on the region of interest specifically in the female strongly suggests it to be the case.

We also report that cRE is specifically enriched for CTCF specifically in female cells. Role of CTCF in *XIST* regulation was ruled out based on the observation that it does not exhibit a differential enrichment between male and female fibroblasts or XIST^+^ and XIST^-^ human ES cells (51). However, the results obtained in our study unequivocally demonstrate it to be specifically enriched on the cRE only on Xi. By performing knockdown and overexpression experiments, we show that XIST is positively regulated by CTCF in addition to YY1. Interestingly, XIST levels increase dramatically when over-expression of YY1 or CTCF is coupled with the knockdown of OCT4 and SOX2 in NT2/D1 cells suggesting that the balance of activators (YY1 and CTCF) and repressors (pluripotency factors) determines *XIST* transcriptional status.

Since overexpression and knockdown based results can be confounded by indirect effects, we made use of CRISPR-dCas9-KRAB based interference approach to specifically target cRE and address its significance more directly. Findings obtained with this strategy corroborated the significance of newly identified cRE in manifesting transcriptional regulation of *XIST* in CTCF and YY1-dependent manner. Therefore, we propose that at the onset of XCI, the cRE is kept repressed by pluripotency factors. However, in the differentiated cells which do not express pluripotency factors, cRE on the Xi is now bound by CTCF **(Figure 6 legend)**. As a result, XIST continues to be synthesized from Xi and is involved in maintaining silencing. Whether the inconsistency in XIST expression observed in various female human ES lines can plausibly be a consequence of the variability in the cRE function would be a promising line of future investigation.

In conclusion, we have identified a novel cRE for *XIST*/*Xist* that is controlled by developmentally regulated factors, OCT4, SOX2 and NANOG as the repressors and the chromatin organizer CTCF/YY1 as activators. Since all the identified regulators are expressed to a similar extent in male and female cells, it is intriguing that they are able to regulate the *XIST* expression in Xi-specific manner. This is suggestive of additional factors/ mechanisms contributing towards such specificity, which deserves further investigation. Determining if this feature is conserved in other eutherians as well will provide useful clues regarding the conserved mechanisms governing *XIST* transcription and provide key insights into evolutionary conservation of the phenomenon of X inactivation.

## MATERIALS AND METHODS

### Cell Culture

Human embryonic carcinoma cell line NT2/D1 (NTERA2-clone D1) was obtained as a kind gift from Dr. Peter Andrews, University of Sheffield, UK. They were grown in Dulbecco’s Modified Eagle’s Medium with sodium pyruvate, high glucose (DMEM, Sigma-Aldrich, St. Louis, Missouri, USA) supplemented with 10% foetal bovine serum (Invitrogen, Carlsbad, California, USA), 2 mM L-glutamine (Invitrogen, Carlsbad, California, USA) and penicillin-streptomycin (Invitrogen, Carlsbad, California, USA) and maintained at 37°C under 5% CO2 atmosphere. NT2/D1 cells were passaged upon reaching 70% confluency by gentle scraping and trypsin was not used for passaging. Human embryonic kidney cells (HEK293T) were grown in DMEM (Invitrogen, Carlsbad, California, USA) without sodium pyruvate, high glucose, supplemented with 10% foetal bovine serum (Invitrogen, Carlsbad, California, USA) and penicillin-streptomycin (Invitrogen, Carlsbad, California, USA) and maintained at 37°C under 5% CO2 atmosphere. HEK293T cells were maintained passaged upon reaching 70-80% confluency using 0.05% trypsin (Invitrogen, Carlsbad, California, USA). DLD1 cell were grown in RPMI (Gibco, Grand Island, NY, USA) supplemented with 10% foetal bovine serum (Invitrogen, Carlsbad, California, USA) and penicillin-streptomycin (Invitrogen, Carlsbad, California, USA) and maintained at 37°C under 5% CO2 atmosphere. DLD1 cells were passaged upon reaching 70-80% confluency using 0.05% trypsin (Invitrogen, Carlsbad, California, USA).

### RA-mediated differentiation of NT2/D1 cells

All-trans-retinoic acid (RA) (procured from Sigma-Aldrich, St. Louis, Missouri, USA) used for inducing differentiation of NT2/D1 cells was reconstituted at a concentration of 5 mg/ml in DMSO (Sigma-Aldrich, St. Louis, Missouri, USA) and stored at −80°C. For differentiation experiments, NT2/D1 cells were harvested using 0.05% trypsin, resuspended in fresh medium and seeded at a density of 0.15 Χ 10^6^ cells in 6-well plate or 1 Χ 10^6^ cells in 100 mm tissue culture dish (Corning, New York, USA). Cells were allowed to grow for 24 hours following which few cells were harvested for RNA and protein extractions or crosslinked for ChIP as 0 day control. RA was added to the remaining wells/plates at a concentration of 13.7 μM for the remaining 6 days. Each day cells were either replenished with fresh medium and RA or harvested as day 1/2/3/4/5/6 samples.

### Molecular Cloning

To characterize *XIST* promoter, genomic regions ranging from +50 bp to −4408 bp upstream of *XIST* TSS were cloned into a promoter-less reporter vector – pGL3 Basic procured from Promega (Wisconsin, Madison, USA). *XIST* genomic regions were PCR amplified from the genomic DNA extracted from NT2/D1cells using specific primer pairs (Supp Table 1) followed by restriction enzyme digestion of the vector and insert and ligation using T4 DNA ligase (New England Biolabs, MA, USA). The positive clones were confirmed by sequencing.

**Table 1.**
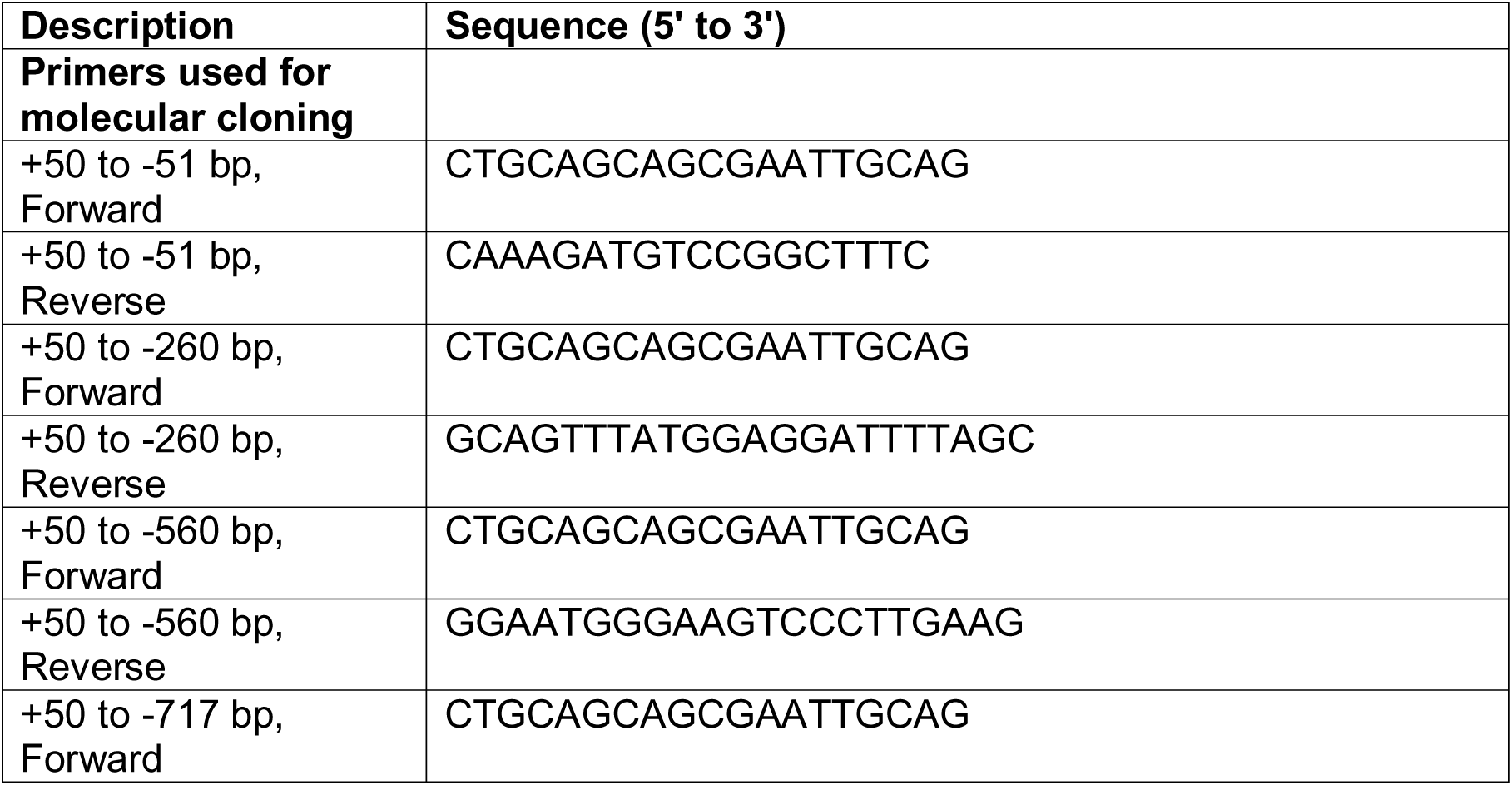
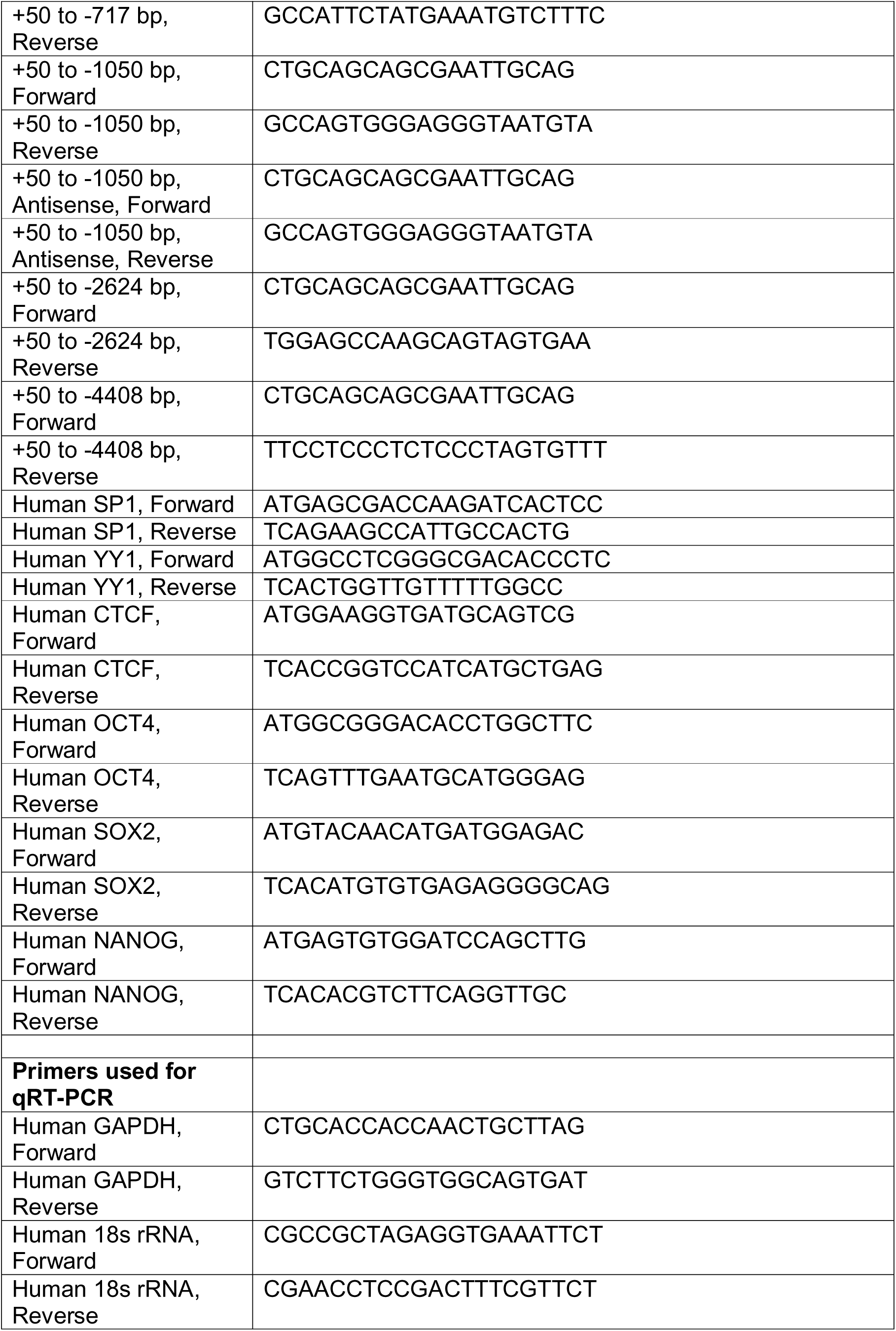
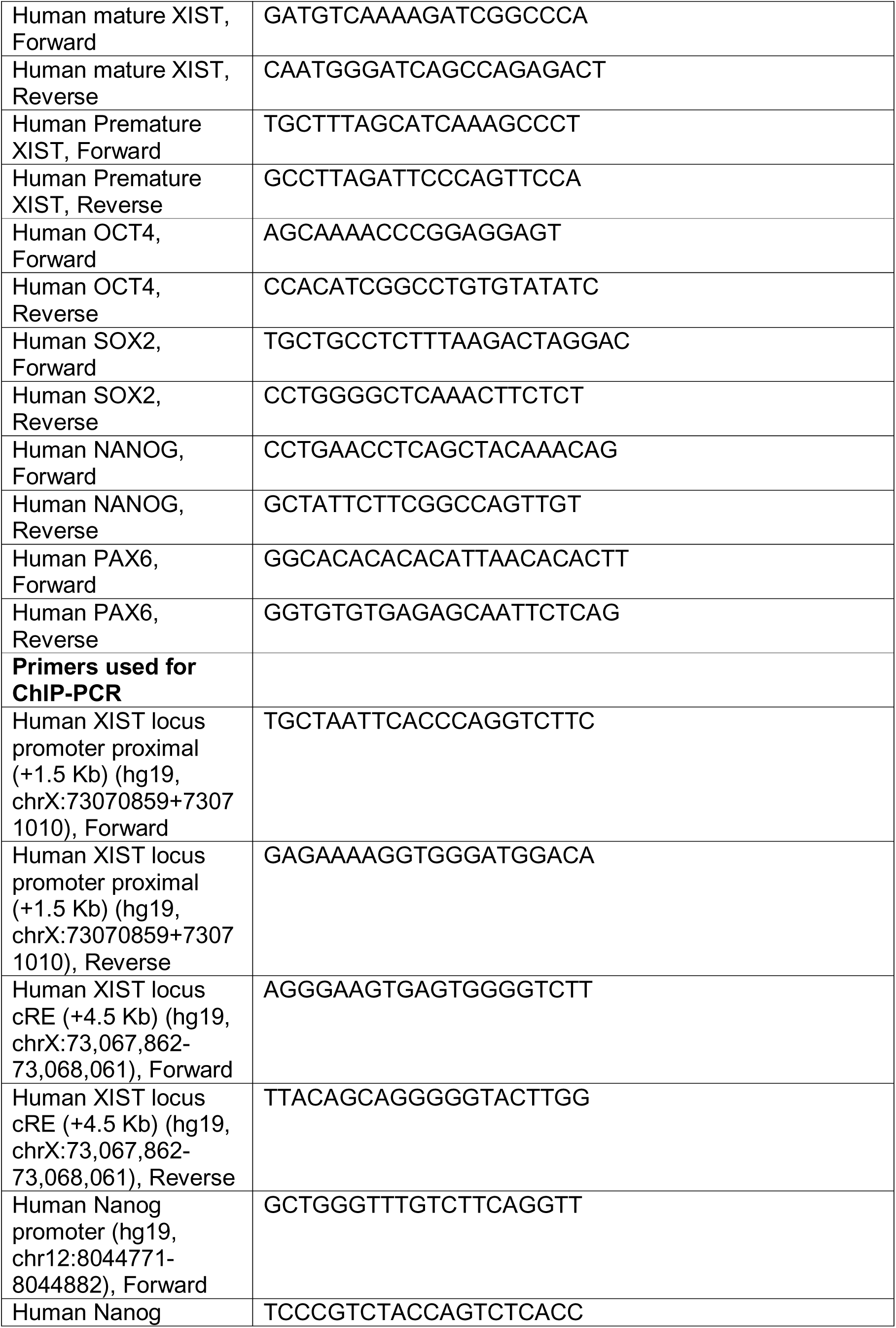
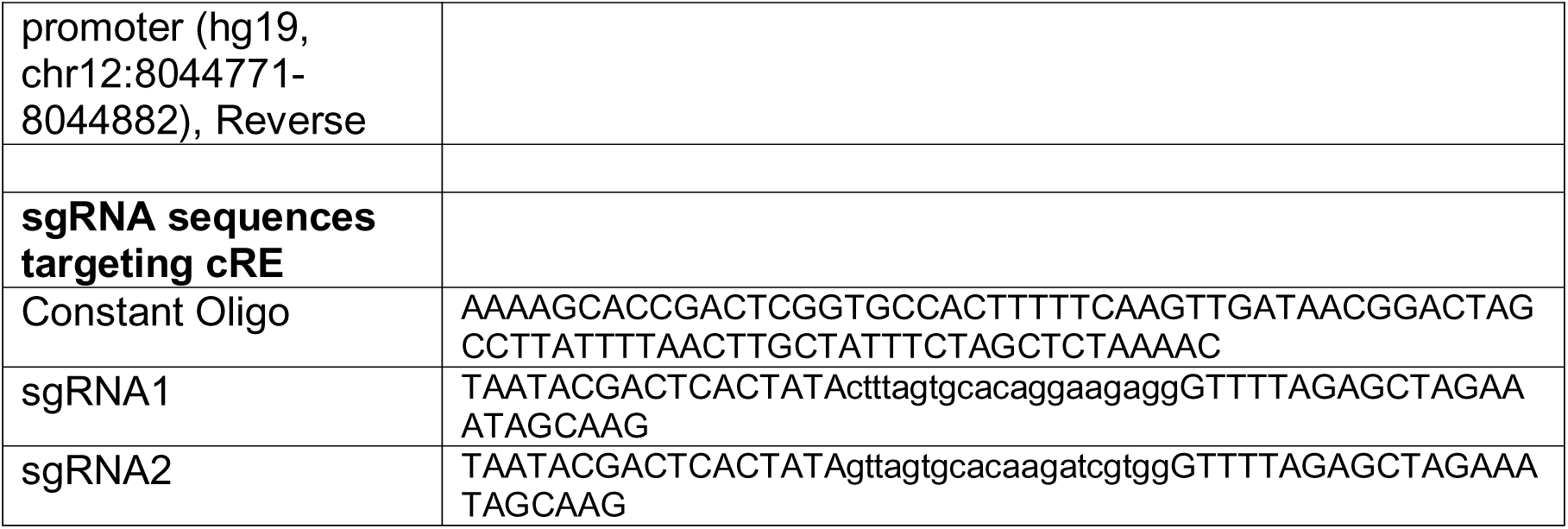
List of oligonucleotide primers used.

### Transfection of DNA, siRNA and gRNA

NT2/D1 or HEK293T cells were transfected with the equimolar of DNA constructs using Lipofectamine 2000 and siRNA using Lipofectamine RNAiMax (Invitrogen, Carlsbad, California, USA) as per manufacturer’s guidelines. HEK293T cells were transfected with plasmid for dCas9-KRAB and gRNA mix using Lipofectamine 3000 as per manufacturer’s guidelines.

### Luciferase reporter Assay

Luciferase reporter assays were performed using the Dual luciferase assay kit from Promega (Madison, Wisconsin, USA). Cells were transfected with Firefly luciferase and Renilla luciferase DNA constructs. After harvesting, the cells were lysed using 1X Passive lysis buffer (Promega, Madison, Wisconsin, USA) as per manufacturer’s instructions. The lysates and substrates were mixed in the optical bottom 96-well plate (ThermoFisher Scientific, Waltham, Massachusetts, United States) according to the guidelines provided by Promega. The reporter activities were measured using luminometry mode on the Varioskan machine (ThermoFisher Scientific, Waltham, Massachusetts, United States). In all the assays, Renilla luciferase activity measurement served as an internal control. The fold change was calculated with respect to either the vector control or 0 day control as and when mentioned.

### RNA extraction and cDNA synthesis

Total RNA was extracted using the Trizol reagent (Invitrogen, Carlsbad, California, USA). 1 to 2 µg of RNA was used for DNase treatment as per the protocol provided by the manufacturer (Promega, Madison, Wisconsin, USA). This was followed by cDNA synthesis using reverse transcriptase kits from Applied Biosystems (High capacity cDNA synthesis kit) (Foster City, California, USA). Additionally, minus RT control was set up to verify the efficiency of DNase treatment. The synthesized cDNA was used to set up quantitative real-time PCR (qRT-PCR).

### Quantitative real-time PCR

Quantitative real time PCR was performed using SYBR green chemistry (Roche) with specific set of primer pairs (Supp Table 1) using ViiA7 thermal cycler (Applied Biosystems). Changes in threshold cycles were calculated by subtracting the Ct values of the gene of interest from that of housekeeping control (for qRT-PCR) [Ct(target genes) – Ct(18s rRNA)]. ΔCt values of specific target genes from the experimental samples were then subtracted from their respective control samples to generate ΔΔCt values. The fold changes were calculated using the formula: 2^(-(ΔΔCt value)). For quantification after ChIP, DNA recovered post ChIP and %Input are used as templates for the PCR with specific set of primers. For quantification of enrichment, the efficiency of chromatin immunoprecipitation of particular genomic locus was calculated as follows :

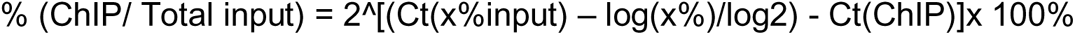

Relative occupancy was calculated as a ratio of specific signal over background: Occupancy= % input (specific loci) / % input (background loci)

### Protein extraction and immunoblotting

Cell pellets were resuspended in RIPA buffer (10 mM Tris (pH 8.0), 1 mM EDTA (pH 8.0), 0.5 mM EGTA, 1% Triton X-100, 0.1% sodium deoxycholate, 0.1% SDS, 140 mM NaCl) containing 1X protease inhibitors (procured from Roche, Basel, Switzerland) and lysed by repeated freeze-thaw cycles. The lysates were centrifuged at 14000 rpm, 4°C, 30 minutes to eliminate the cellular debris. The supernatant was collected in the fresh microfuge tube. The concentrations of protein were determined by performing BCA assay (purchased from ThermoFisher Scientific, Waltham, MA, USA). Equal amounts of protein lysates were boiled in 1X Laemmli buffer (0.5 M Tris-HCl pH 6.8, 28% glycerol, 9% SDS, 5% 2-mercaptoethanol, 0.01% bromophenol blue) for 10-15 minutes and subjected to electrophoresis on a polyacrylamide gel. The separated proteins were transferred onto PVDF membrane (Millipore, Billerica, Massachusetts, USA) using phosphate based transfer buffer (10 mM sodium phosphate monobasic, 10 mM sodium phosphate dibasic) at 4°C, 600 mA, 2 hours. After the completion of transfer, membranes were blocked in 5% skimmed milk, incubated overnight at 4°C with the primary antibodies prepared in 5% BSA. The membranes were washed thrice with the buffer containing 20 mM Tris buffer pH 7.4, 500 mM NaCl and 0.1% tween 20 (TST) the next day and incubated with the appropriate secondary antibodies conjugated with horseradish peroxidase for an hour at room temperature. Following this, the membranes were again washed thrice with TST buffer. The blots were developed using Immobilon Western Chemiluminescent HRP Substrate (Millipore, Billerica, MA, USA) and detected using ImageQuant LAS 4000 (GE Healthcare, Piscataway, NJ, USA) according to the manufacturer’s instructions.

### Chromatin immunoprecipitation

ChIP was performed as per X-ChIP protocol. Briefly, 10 µg of sonicated chromatin (average length 150-400 bp) was incubated with 1 µg of specific antibody and incubated on an end-to-end rocker at 4°C overnight. Each ChIP was performed at least 3 times as indicated in the figure legends and quantitative real-time PCR was set up. Primers used for the ChIP-qPCR analysis are listed in **Table 1**.

### Sequential ChIP assay

The method followed for the sequential ChIP assay was same as that described above. ChIP was started with 200 µg of chromatin and the 1^st^ elution was done using the buffer containing 0.1 M NaHCO_3_, 1% SDS and 10 mM DTT. The eluate was subjected to second ChIP with either Rabbit IgG or CTCF antibodies.

### RNA FISH

The RNA FISH protocol followed was as described by Chaumeil et al. 2005. Briefly, the probe for RNA FISH was prepared with 2 µg of BAC RP183A17 using Nick translation kit (10976776001) purchased from Roche (Basel, Switzerland). Salmon sperm DNA and cot1 DNA were added to the probe to mask the repetitive sequence. Prior to hybridizations, the cells on the coverslips were washed with RNase-free PBS followed by permeabilization using freshly made Cystoskeletal buffer [10 mM PIPES (pH 7.0), 100 mM NaCl, 300 mM Sucrose, 3 mM MgCl_2_] containing 0.5% TritonX-100 for 4 to 5 minutes on ice. Nuclei are fixed using 4% paraformaldehyde for 10 minutes at room temperature. The coverslips were washed twice with 70% ethanol and then subjected to dehydration by sequentially incubating the coverslips in 80%, 95% and 100% ethanol for 3 minutes each, followed by air drying. The nuclei were then subjected to hybridization step in the dark and humidified chamber at 37°C for 24 hours. After incubation, the coverslips are washed thrice with 50% formamide + 2X SSC buffer (pH 7.2) for 5 minutes each at 42°C, followed by washing thrice with 2X SSC buffer for 5 minutes each at 42°C. Counterstained with DAPI and washed twice with 2X SSC buffer. The coverslips were then mounted on a clean slide for microscopic observation.

### Metaphase spread preparation

A semiconfluent culture of NT2D1 cells (∼60% confluency), was subjected to metaphase arrest by treating these cells with 0.1 µg/ml colcemid for 90 min at 37°C. Cells were harvested by trypsinization, followed by hypotonic treatment (0.075 M KCl) for 30 min at 37°C, which was terminated by fixing cells with 4–5 drops of chilled fixative solution (Methanol: Acetic Acid, 3:1), followed by centrifugation at 1000 rpm for 10 min at 4°C. The cell pellet was washed three times with fixative solution to remove cell debris and resuspended in fresh fixative solution, based on the amount of cell pellet recovered. This suspension was dropped from a height onto a clean glass slide and air-dried.

### 2-Dimensional in situ hybridization

Whole chromosome paint for human X-chromosome (WCP-X) was obtained from Applied Spectral Imaging (ASI), Israel. WCP-X (5 μl) was equilibrated at 37°C for 5 minutes, denatured at 80°C for 5 min, and quick chilled on ice for 2 minutes followed by a pre-annealing at 37°C for 30 minutes. The denatured probe (3-4 μl) was spotted onto a pre-marked area of a glass slide containing fixed interphase nuclei and metaphases (as prepared above) and covered with an 18mm X 18mm coverslip, sealed with nail polish and co-denatured at 72-74°C for 1.5 minutes and hybridized for 36-48 hours in a humidified box at 37°C. Post hybridization, the coverslip was removed carefully and the glass slides were washed in 50% FA/2XSSC (pH 7.4), thrice for 5 minutes each at 45°C, followed by three washes for 5 minutes each in 1XSSC at 45°C with gentle agitation. Slides were briefly dipped in 0.1% Tween20/4X SSC, and counterstained with 4’,6-Diamidino-2-Phenylindole (DAPI) for 2 minutes, washed in 2X SSC and mounted in Antifade and stored at 4°C until they were imaged. Metaphases and interphase nuclei were imaged under an Axioimager Z2 immunofluorescence microscope (Zeiss) (63X Objective, N.A 1.4).

### gRNA generation for dCas9-KRAB based repression of cRE

Two sgRNA for cRE were designed using CHOPCHOP webtool (73). For each sgRNA, a 60 base oligonucleotide (“gene-specific oligo”) containing (1) a promoter for in vitro transcription, (2) the 20 base spacer region specific to the target site and (3) an overlap region that anneals to the constant oligonucleotide was synthesized. sgRNA were generated by *in vitro* transcription (IVT). In short, primer carrying T7 promoter site and sgRNA sequence was annealed with the constant oligo and used for IVT. IVT performed with MAXIscript™ T7 Transcription Kit (Thermofisher) using protocol provided with the kit.

### Antibodies, siRNAs and other reagents

SP1 (5931S) antibody for immunoblotting were procured from Cell Signaling Technology (Danvers, USA), YY1 (ab12132) (sc-7341) antibodies for western blotting was purchased from Abcam, SOX2 (AF2018) and NANOG (AF1997) antibodies for immunoblotting and ChIP were purchased from R & D (Menomonie, USA), OCT4 antibody (sc-9081) for western blotting and ChIP and goat-HRP secondary antibody were purchased from Santacruz Biotechnologies (Dallas, Texas, USA), CTCF antibody (07–729) procured from Millipore/Upstate (Billerica, Massachusetts, USA) was used for all of the ChIP experiments. CTCF antibody (sc-21298) purchased from Santacruz Biotechnologies (Dallas, Texas, USA) was used for immunoblotting, Normal rabbit IgG (12–370) and Normal mouse IgG (12–371) for ChIP were purchased from Millipore (Billerica, Massachusetts, USA), β-ACTIN (VMA00048) and γ-TUBULIN primary antibodies, mouse-HRP and rabbit-HRP secondary antibodies were purchased from BioRad Laboratories (Hercules, California, USA). Histone modification antibodies used for the ChIP were as follows - H3K4me (Millipore, 07-436), H3K27ac (Abcam, ab4729), H3K27me3 (Abcam, ab6002), H3K4me3 (Millipore, 17-614) and H3K9me3 (Millipore, 07-523) The following antibodies were used for ChIP-qPCRs following dCas9-KRAB targeting: anti-FLAG (Sigma: F-3165), anti-CTCF (Abcam: 612148), anti-H3K9me3 (Millipore 07-442), anti-YY1 (SantaCruz: sc-7341).

### Sequencing analysis for ChIP-Seq

SRR/DRR/ERR IDs of the datasets used are provided. Datasets used tabulated by cell line, the files can be downloaded from the link (https://www.ebi.ac.uk/ena) by providing the IDs given below as search terms.

H9 cells: OCT4 (SRR5642847), Nanog (SRR5642845); HEK293 cells: CTCF (DRR014670); DLD1 cells: CTCF (DRR014660); Female breast epithelium cells (ENCODE): CTCF (SRR6213076, Input SRR6213541); Male breast epithelium cells (ENCODE): CTCF (SRR6213724, Input SRR6214303); MCF7: CTCF (SRR577680,SRR577679) (ENCODE), Hela: CTCF (SRR227659, SRR227660) NT2/D1: OCT4 (SRR1640282) (This dataset is generated on the platform SOLiD3.0).

All Illumina datasets were aligned with Bowtie 2 (71) to the hg19 reference assembly. Following which MACS2 (72) was used to identify significant peaks relative to control where available. The p value cut-off was set at 0.05, MACS2 callpeak uses the *t* test to determine significance of the enrichment at a certain location over control. The resulting .narrowPeak files were used to plot the figures regarding ChIP Seq peaks using the GViz package in Rstudio. SOLiD 3.0 dataset for the NT2/D1 OCT4 ChIP-Seq was aligned to the hg19 reference genome using Bowtie1 following which the downstream processing was as indicated above.

## DATA AVAILABILITY

Files for the datasets used for analysis can be downloaded from the link (https://www.ebi.ac.uk/ena).

## ACKNOWLEDGEMENTS

Authors wish to thank Dr. Peter Andrews (University of Sheffield) for kindly gifting NT2/D1 cells; Dr. Ajay Labade and Madhu Kabra (IISER Pune) for providing help with the FISH experiments; Dr. Mukul Rawat and Sneha Tripathi (IISER Pune) for helping with the imaging and analysis. Figure 5D was created using **Biorender**.com.

## AUTHOR CONTRIBUTIONS

RS and SG conceived the project and designed experiments. RS performed all experiments except X chromosome paint (Fig 1E) and dCas9-KRAB based targeting (Figure 5D-G), interpreted data and wrote the manuscript. AS performed and analyzed dCas9-KRAB experiments in HEK293T cells. AK performed bioinformatics analysis and interpretations and also helped with acquisition of the RNA FISH images. KS performed X chromosome paint. SG interpreted data, supervised the project and wrote the manuscript. All authors read and approved the final manuscript.

